# An interpretable meta-clustering framework for single-cell RNA-Seq data integration and evaluation

**DOI:** 10.1101/2021.03.29.437525

**Authors:** Zhiyuan Hu, Ahmed A. Ahmed, Christopher Yau

## Abstract

Single-cell RNA sequencing (scRNA-Seq) datasets that are produced from clinical samples are often confounded by batch effects and inter-patient variability. Existing batch effect removal methods typically require strong assumptions on the composition of cell populations being near identical across patients. Here we present a novel meta-clustering workflow, CIDER, based on inter-group similarity measures. We demonstrate that CIDER outperforms other scRNA-Seq clustering methods and integration approaches in both simulated and real datasets. Moreover, we show that CIDER can be used to assess the biological correctness of integration in real datasets, while it does not require the existence of prior cellular annotations.

## Introduction

The widespread adoption of single-cell RNA sequencing (scRNA-Seq) as a modality for the investigation of functional cellular heterogeneity means it is now routine for multiple datasets to be generated from the same type of tissues and organs across a number of individuals. Integration of multiple scRNA-seq datasets can provide more comprehensive interpretations by borrowing information across experiments and even species^1^. However, the data from multiple experiments are often confounded by inter-batch or inter-donor variability.

Existing clustering workflows can effectively identify cell populations in batch-effect-free datasets^2^, by partitioning cells based on the inter-cell distance matrix computed from the expression data of high variance genes (HVGs) or the derived principal components. For example, SC3 constructs the distance matrix by applying Euclidean, Pearson and Spearman metrics on the expression data of HVGs and transfers this distance matrix by principal component analysis (PCA) or graph Laplacian transformation, before consensus clustering^3^. RaceID computes the distance matrix in the same way as SC3 but provides more options of distance measures, including Kendall and proportionality^4^. Seurat v3 calculates Euclidean distances from the principal components and then infers the graph of shared nearest neighbors for the subsequent graph-based clustering, such as Louvain clustering^5^. However, distance measurements used by these workflows cannot effectively distinguish biological variation from the technical one and, thus, their performance is compromised in datasets confounded by batch effects or other variability caused by unwanted or unexplained factors.

In data confounded by batch effects, workflows combining batch correction or integration methods and downstream clustering algorithms are used to identify cell populations. Some existing batch correction and integration methods can efficiently correct the gene expression or dimensionality reduction spaces for visualization and other downstream analyses. For example, mutual nearest neighbors (MNN)^6^ uses the cells pairs that are mutually nearest neighbors to compute a vector that aligns multiple batches into a common space. Scanorama^7^ also used the concept of MNNs to merge datasets with substantial improvement in the MNN search strategies. Seurat^8^ exploits canonical correlation analysis (CCA) to compute a subspace and then used the identified MNNs, i.e. “anchors”, to correct the data. Harmony^9^ iteratively diminishes batch effects in the PCA space by soft clustering cells across batches and then adjusting cell positions based on the global and dataset-specific cluster centroids.

However, for the majority of integration methods, performance can vary substantially across data types and scenarios^10^. An additional limitation of the commonly used integration algorithms, e.g., CCA and Harmony, is that they work on the low-dimensional representation, which can be affected by the bias in the initial selection of HVGs. Further, it is often difficult to determine why existing methods drives cells from different batches into the same cluster. This lack of explainability or interpretability can make it difficult to ascertain if integration has been successful.

To address this limitation, we recently introduced the use of meta-clustering to partition scRNA-seq data from ovarian cancer fallopian tube epithelial cells confounded by structured batch effects and inter-patient variability^11^. This method was based on a functional hypothesis that cells from the same biological population (either cell type, subtype or state) share a similar differential expression pattern, i.e., the differentially expressed genes (DEGs) having more weights to determine cell classes compared to other genes. Moreover, these DEGs are less affected by batch effects by regressing out the unwanted factors. In this work, we present a scalable version of this methodology, and demonstrate its generalizable utility for wider application.

Here we introduce a novel similarity metric based on Inter-group Differential ExpRession (IDER) and propose a workflow of Clustering by IDER (CIDER). We demonstrate that the performance of CIDER is comparable or superior to existing clustering workflows applied on uncorrected and batch-corrected datasets in a variety of scenarios for both simulated and real scRNA-seq data. Since the distance measurement of CIDER uses a different principle from other measurements used in popular integration algorithms, we show that CIDER can also be used as a ground-truth-free evaluation metric for accurately identifying falsely integrated populations.

## Results

### Design of CIDER and proof-of-concept experiment

The core of CIDER is the IDER metric, which can be used to compute the similarity between two groups of cells across datasets (Fig. 1a). IDER first identifies the differentially expressed signature (DES) for each group of cells against all other cells within the same batch or dataset. Next, a similarity measure is computed by using the consistency of DESs between two groups across datasets. Differential expression in IDER is computed using limma-voom^12^ or limma-trend^13^ which was chosen from a collection of approaches for differentially expression analysis based on a number of performance criteria^14^ (Supplementary Fig. 1a, b).

**Figure 1.**
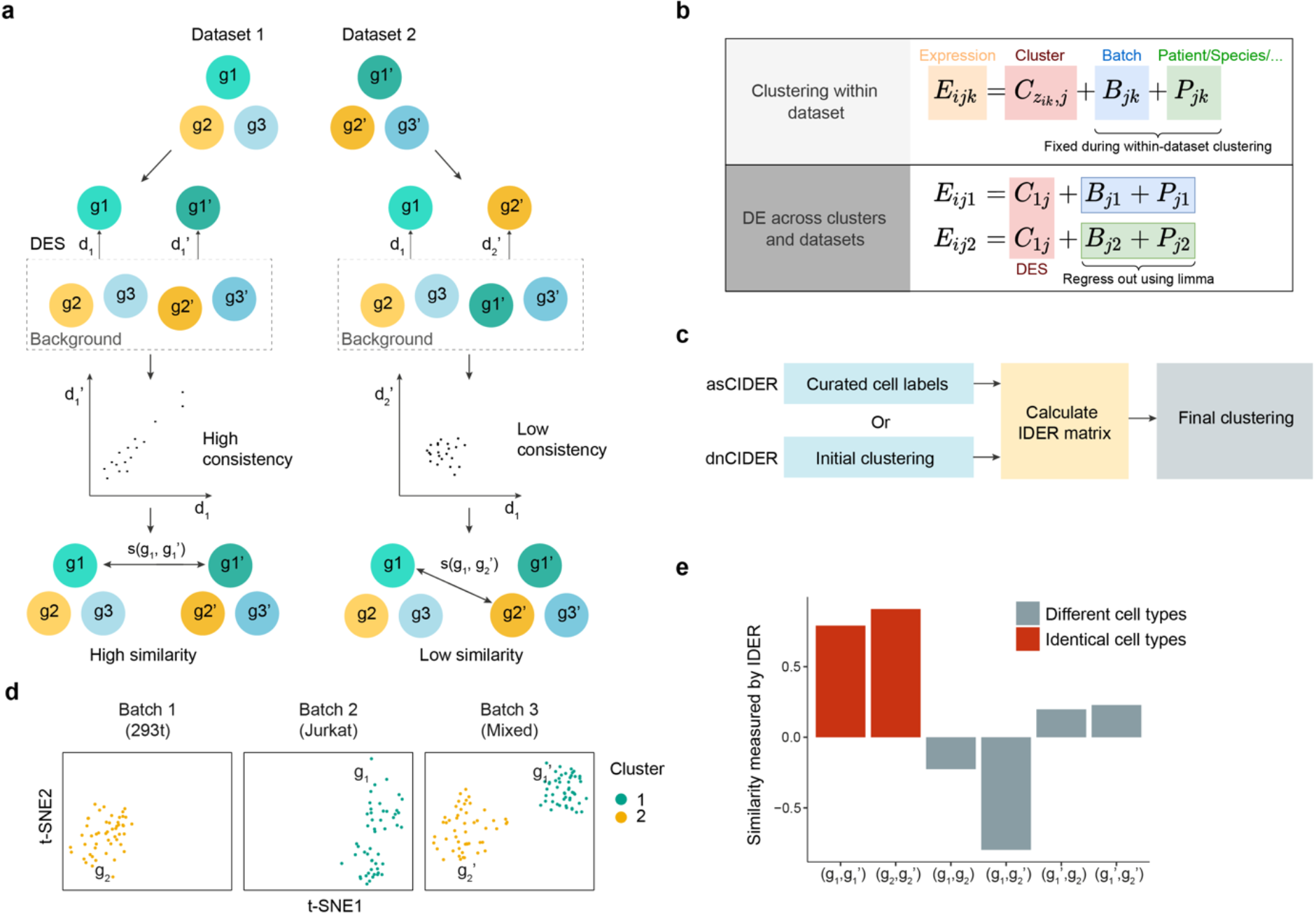
IDEr metric accurately measures the biological similarity between cell groups. **(a)** Schematic diagram shows how the IDER metric measures the inter-group similarity. **(b)** Diagram shows the theoretical justification for CIDER. *E_ijk_* denotes the expression level of gene *j* in cell *i* of batch *k, z_ik_* cell assignment of cells of cell *i*, *C* cluster effect, *B* batch effect and *P* patient effect. In the lower panel, *z_ik_* is set to 1. **(c)** Schematic diagram of asCIDER and dnCIDER. **(d)** t-SNE plots show the cells from three batches of Dataset 1. Each subpanel represents a batch. Cells are colored by the population. Each batch-specific cluster is denoted by a label. **(e)** The IDER metric generated higher similarity between group pairs, (g_1_, g_1_’) and (g_2_, g_2_’), from identical cell types and lower similarity between groups pairs from different cell types.

CIDER is established on the hypothesis that the expression level contains the linear combination of the effects of cluster, batch and patient (Fig. 1b). The within-dataset clustering enables the identification of the cluster effect (i.e., cell assignment) for a given dataset, as the confounded effect (e.g., batch effects, inter-patient variability or inter-species variability) is a constant within the same dataset. Once the cell assignments are completed for all datasets, we use limma to regress out the confounded effects across datasets and identify consistent cluster effects, represented by DESs, from multiple datasets. Groups with the consistent cluster effect will be merged into one final cluster. In the workflow of CIDER, IDER is used to measure the pairwise inter-group similarity among the batch-specific initial clusters (Fig. 1c). These initial clusters can be either curated annotations or clustering results. The output of the IDER step, i.e., a similarity matrix, is used to merge the connected initial clusters into final cross-batch clusters. Depending on how the initial clusters were derived, we named the CIDER workflows as *de novo* CIDER (dnCIDER), where initial clusters were the output of a clustering algorithm, and assisted CIDER (asCIDER), where initial clusters were curated annotations of cell populations. These two scenarios were considered in our benchmarking because they are common in the real-world usage.

We set about to test if the IDER metric could accurately estimate the cluster effects and regress out confounded ones in data confounded by batch effects. As a proof-of-concept experiment, we applied it to a multiple cell line dataset^15^ (Dataset 1), in which there were three batches corresponding to pure 293T cells, pure Jurkat cells and a 50/50 mixture of both cell lines. The IDER metric was used to calculate the pairwise similarity among four groups from these three batches (Fig. 1d & Supplementary Fig. 2). We showed that the similarity computed by IDER was higher for the group pairs from the identical cell type compared to the pairs from different cell types (Fig. 1e), demonstrating the utility of IDER as a metric to identify cluster similarity across datasets when confounded by batch effects.

### Benchmarking on simulated and real data

To test the accuracy of identifying populations, we benchmarked CIDER against three singlecell clustering approaches (Seurat v3-Louvain^5^, SC3^3^ and RaceID^4^) and four workflows that combined integration approaches (Seurat-CCA^8^, MNN^6^, Scanorama^7^ and Harmony^9^) and clustering. We used a simulated dataset (Dataset 2, Supplementary Fig. S2, Supplementary Table 1), where three batches composed of non-identical compositions of populations (Supplementary Fig. 3a). In this scenario, the cross-batch similarity computed by CIDER correctly recognized the connection among initial clusters (Fig. 2a, b). In contrast, MNN and CCA overcorrected the batch effects, leading to the incorrect merging of disparate populations as previously reported^7^ (Supplementary Fig. 3b, c). To quantitatively compare their performance, we computed the adjusted Rand indexes (ARIs) between cell labels and clustering results (ARI_popuiation_) or the ARIs between batches and clustering results (ARI_batch_). The experiment replicates (n = 20) confirmed that CIDER robustly outperformed MNN and CCA in this scenario of non-identical cellular compositions (Fig. 2b).

**Figure 2.**
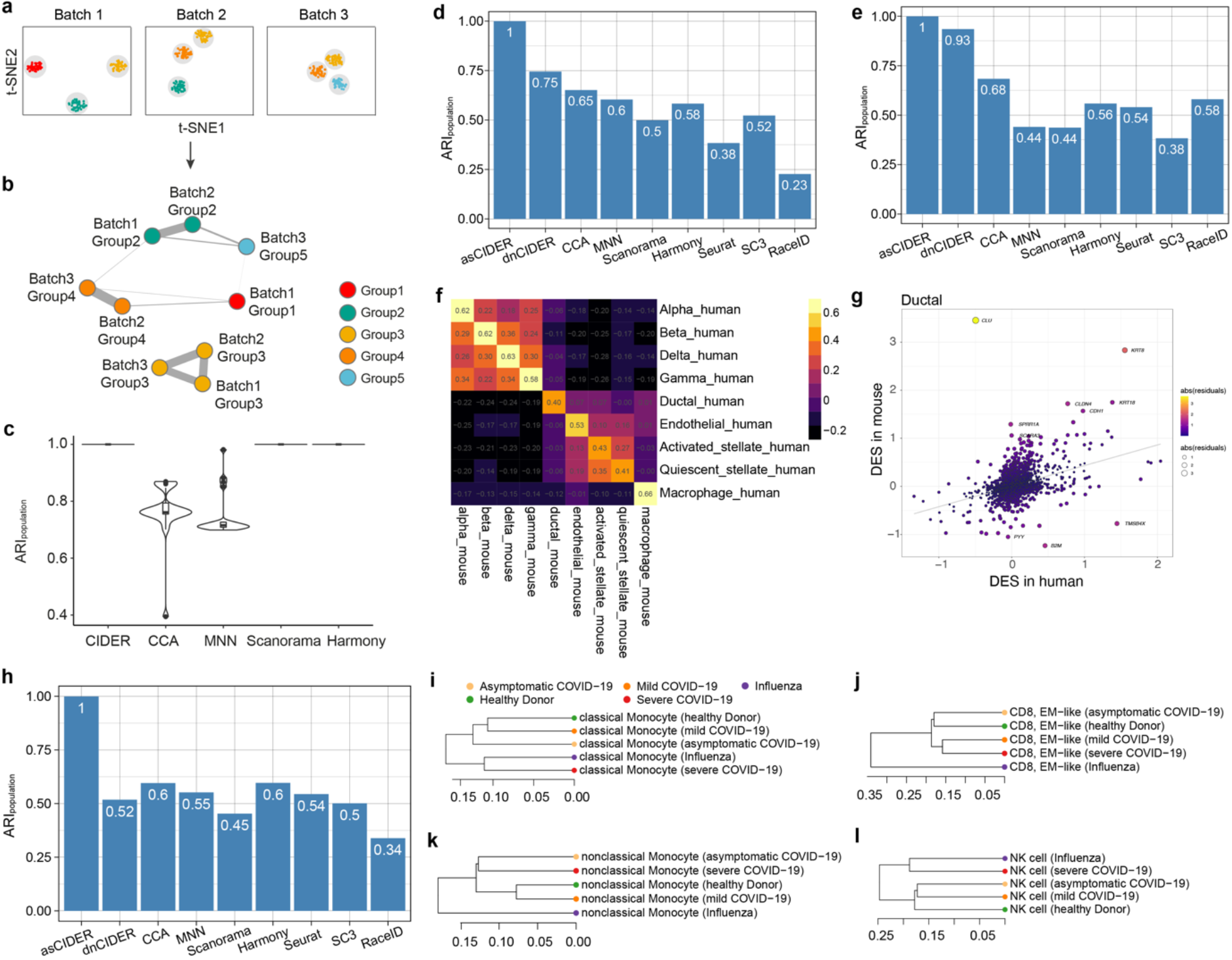
CIDER accurately identifies cross-batch populations. **(a)** t-SNE plots show the nine initial clusters from three batches of the simulated dataset (Dataset 2). Cells are colored by populations. **(b)** The graph network shows the similarity among initial clusters. Vertexes represent initial clusters, colored by populations. The width of edges represents the similarity levels. **(c)** Distribution of ARI_popuiation_ for 20 replicates of simulated data across integration workflows and clustering algorithms. The x-axis denotes clustering and integration workflows. The y-axis denotes the workflow performance by calculating ARIs between cell populations and clustering results. **(d-e)** Distribution of ARI_popuiation_ across integration workflows and clustering algorithms for the human PBMC data (Dataset 3) (d) the cross-species pancreas data (Dataset 4) (e). **(f)** Heatmap shows the inter-group similarity between mouse populations (x-axis) and human ones (y-axis). Cells are colored by the similarity levels, as shown by the numbers. **(g)** Scatter plot shows genes driving the similarity and dissimilarity between human ductal cells and mouse ones. The x- and y-axes denote the DESs in human and mice. Each dot is a gene, colored and sized by the residuals. The grey line is the linear regression line. **(h)** The distribution of ARI_popuiation_ of benchmarked algorithms for Dataset 5. **(i-l)** Dendrograms show the local relationships of the classical monocyte population (i), the effector memory (EM)-like CD8 T cell population (j), and nonclassical monocyte population (k) and the natural killer (NK) population (l) from patients with different conditions.

We next tested CIDER with data from human peripheral blood mononuclear cells (PBMCs; Dataset 3; Supplementary Fig. 4a, b)^15^. Both dnCIDER and asCIDER outperformed other batch-correction and clustering workflows regarding the accuracy of identifying populations (Fig. 2d, Supplementary Fig. 4c, d). Apart from partitioning cells into clusters, CIDER also revealed the underlying structure among cell populations. For example, DESs between two monocyte subtypes or between CD4 and CD8 T cells showed relatively high similarity (Supplementary Fig. 4e).

Given the increasing requirement of cross-species comparative analysis, we benchmarked CIDER on pancreatic data that contains both human and mouse data (Dataset 4; Supplementary Fig. 5a, b)^16^. CIDER workflows outperformed other pipelines with respect to ARI_popuiation_, and the capability to correct batch effects was comparable to Seurat-CCA and slightly better than MNN and Scanorama (Fig. 2e, Supplementary Fig. 5c). Computational speed was comparable to Seurat-CCA, MNN and much faster than SC3 and RaceID. We found that for different cell types, the cross-species agreement of DESs was inconsistent (Fig. 5f). We examined the genes that drove the agreement or against it, enabled by the interpretability features of CIDER, and found that a lack of high-agreement genes compromised the cross-species similarity of the ductal population (Fig. 2g), compared to the alpha population (Supplementary Fig. 6).

We next tested on the recent PBMC data (Dataset 5)^17^ collected from healthy donors, patients with COVID-19 and patients with influenza, where the health conditions of the donors were regarded as a batch. Albeit that Seurat-CCA was applied in the original study, the low ARI_batch_ of Seurat, SC3 and RaceID clustering results suggested that this dataset was mildly confounded by batch effects (Supplementary Fig. 7a). Among the benchmarked methods, asCIDER had the highest ARI_popuiation_, while the other methods except RaceID had similar ARI_popuiation_ values between 0.45 and 0.60 (Fig. 2g). This was likely due to the manually merged cell annotations ^17^, where the similarity between defined cell populations might not reflect the statistical similarity defined by these clustering and integration algorithms. The CIDER methods consumed comparable runtime with Seurat-CCA, SC3 and RaceID (Supplementary Fig. 7b). Moreover, the inter-group distance matrix generated by asCIDER unraveled the local relationship of a certain cell population from various conditions. For example, the populations of classical monocytes, natural killer (NK) cells, red blood cells (RBC), dendritic cells (DC) and lgG+ B cells from patients with severe COVID-19 were more akin to the ones from patients with Influenza than the ones from patients with mild or asymptomatic COVID-19, while nonclassical monocytes and effector memory (EM)-like CD8 T cells were not (Fig. 2h-k, Supplementary Fig. 7c-e). Overall, it suggests that asCIDER and dnCIDER not only performed well on data confounded by technical effects, species difference and disease variability, but also provided insights into the relationships across different conditions for a certain cell population.

### Ground-truth-free validation metrics of integration

To benchmark the performance of integration methods, a number of evaluation metrics has been proposed. Most of the assessment metrics for the biological classification require a ground truth, i.e., the curated cell annotations. For example, the local inverse Simpson Index (LISI) represents the population diversity within the neighbors of a given cell by using the lowdimensional representation and cell annotations^9^. Here LISI was used to measure the local diversity of cell population. Another approach is to partition the corrected expression data into clusters and then to calculate the ARI between the clustering result and the prior annotations of cell populations^10^. However, there are relatively few datasets with well-curated annotations of cell populations.

To address this limitation, we embedded CIDER into a workflow of evaluating integration, which does not require the existence of ground truth cell type annotations. After integration by a certain tool, the corrected space can be used to generate cross-batch clusters. The evaluation workflow of CIDER is aimed to estimate the probability of whether cells from a cross-batch cluster were from the same biological population (Fig. 3a).

**Figure 3.**
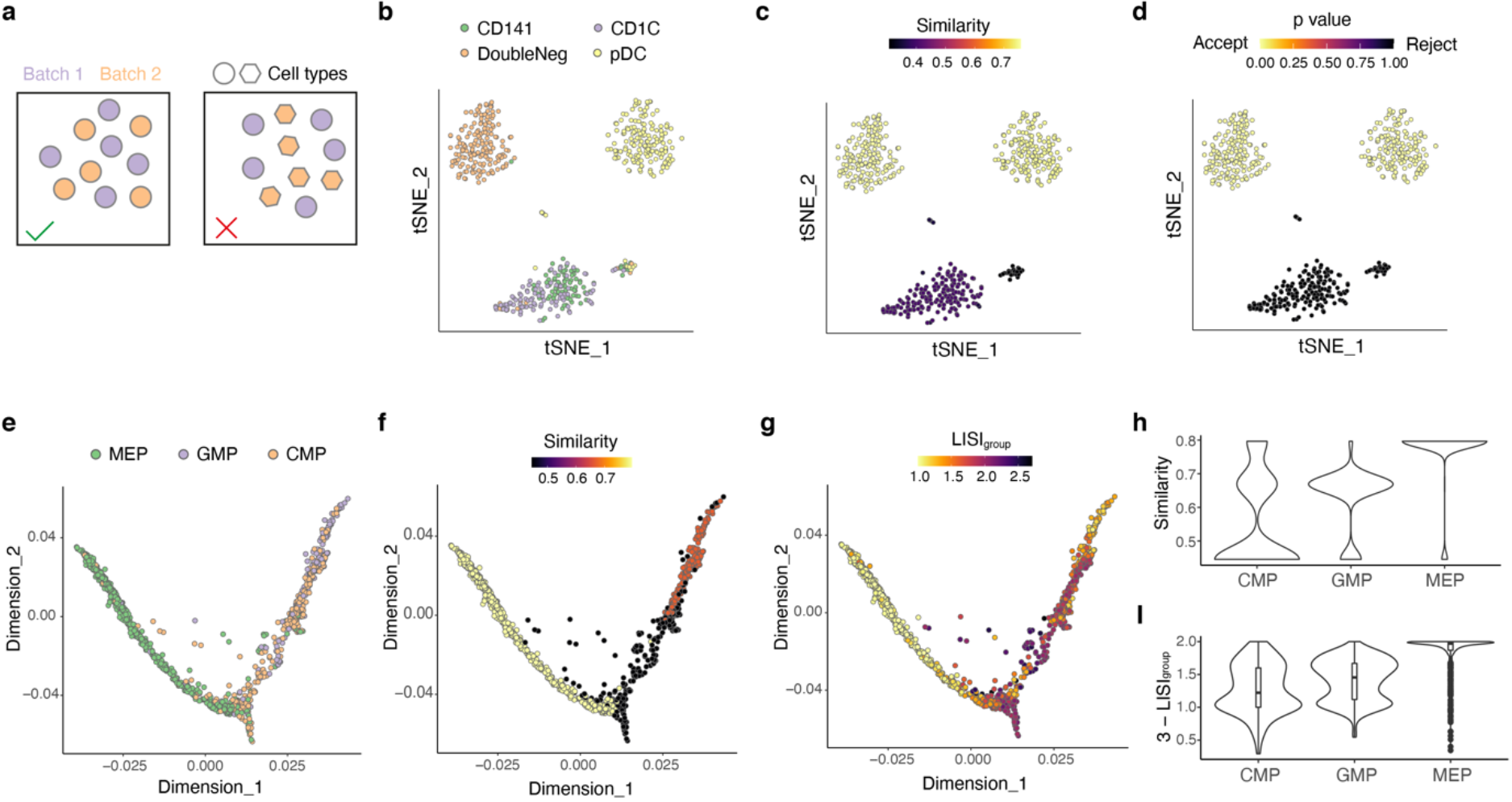
CIDER can evaluate the biological accuracy of integration results without reliance on ground truth. (a) The diagram shows two scenarios, one, where the same cell type from two batches is aligned, and one, where the two different cell types from two batches are falsely aligned. CIDER can identify the second scenario without prior information of cell types. **(b-d)** t-SNE plots of CCA-corrected dendritic cell data (Dataset 5), where cells are colored by cell populations **(b)**, the similarity scores computed by CIDER **(c)** and the empirical p-values computed from the background distribution **(d)**. **(e-g)** Diffusion maps of CCA-corrected mouse hematopoietic progenitor data (Dataset 6), where cells are colored by cell populations **(e)**, the similarity scores computed by CIDER **(f)** and the local purity represented by 3 - LISI_group_ **(g)**. **(h-i)** The distribution of agreement between two experiments for three cell populations, CMP, GMP and MEP, calculated by CIDER **(h)** and LISI **(i)**.

We applied Seurat-CCA on the dendritic cell data^18^ (Dataset 6), where the integration algorithm is prone to merging CD141 cell population and CD1C population incorrectly^10^ (Fig. 3b). By applying CIDER on the corrected low-dimensionality representation, the mixture cluster of the CD141 and CD1C populations was found to have lower similarity than the other two properly aligned clusters: DoubleNeg and pDC (Fig. 3c). The empirical p-value calculated based on a background distribution further confirmed that the mixed cluster should be rejected for alignment (Fig. 3d and Supplementary Fig. 8a-d). The results of CIDER were consistent with LISI (Supplementary Fig. 8e), while the application of LISI requires the correct labels of cell populations.

We next tested CIDER on the mouse hematopoietic progenitor data (Dataset 7)^19,20^ with the continuous data structure. Cells of this dataset were assigned to three populations, the common myeloid progenitor (CMP), the megakaryocyte/erythrocyte progenitor (MEP) and the granulocyte/macrophage progenitor (GMP), and profiled by two platforms, MARS-seq^20^ and Smart-seq2^19^ (Fig. 3e). After integration and dimensionality reduction, the cells around the bifurcating point showed lower levels of agreement between two experiments (Fig. 3f). Based on the ground truth (cell annotations), the results of LISI_group_ also suggested that multiple populations were mixed around the bifurcating point (LISI_group_ ≥ 2; Fig. 3g). The results of CIDER showed that the alignment scores of CMP, the direct ancestry of both MEP and GMP, are lower than those of MEP and GMP between two experiments, which is consistent with the results of LISI (Fig. 3h, i). This is likely due to the higher level of heterogeneity in the predefined CMP population compared to MEP. Taken together, we demonstrated that CIDER was capable of evaluating the local biological similarity without using the predefined cell annotations.

## Discussion

In this work, we presented a meta-clustering framework, CIDER, for scRNA-Seq data integration and evaluation. The benchmarking demonstrated the performance of CIDER regarding the accuracy of recognizing cellular populations, the effectiveness of removing batch effects and the computational complexity.

CIDER used a novel and intuitive strategy that measures the similarity by performing group-level calculations, which stabilize the gene-wise variability. Compared to other distance measures or anchors used for clustering and integration^6-8^, IDER is more interpretable and versatile to quantify biological similarity. CIDER can be exploited for preliminary analysis, standalone clustering or independent validation. Since IDER is built on a different rationale from conventional integration approaches, it can provide alternative insights. Moreover, CIDER can compute a similarity score between cell groups from two conditions, enabling the inference of local relationships based on the expression profiles. Among other methods, Scanorama^7^ can also calculate an alignment score for pairs of datasets for better interpretability, but it is derived from the membership of shared nearest neighbors rather than directly estimated from expression profiles.

CIDER is currently designed for scRNA-Seq data and cannot be used for the integration of single-cell multi-modal data^21,22^. Future work can be focused on adapting the linear model embedded in CIDER for this purpose. Moreover, CIDER is a coarse-grained method based on group-level similarity, while it can be applied to data with continuous structures as we demonstrated.

Taken together, CIDER provides a clustering framework for integrative analysis of multiple scRNA-Seq datasets, enabling identifying cell assignments across datasets and validating the integration output for the assembly of multiple scRNA-Seq datasets.

## Methods

### Measurement of inter-group similarity

The infrastructure of IDER was built on limma-trend^13^ or voom^12^. Both limma-trend and voom estimate the mean-variance relationship non-parametrically by locally weighted regression and then leverage the estimation for DE analysis. The difference between limma-trend and voom is that the mean-variance relationships exploited by them is at the genes level and at the level of individual observations respectively.

Limma methods were selected out of a collection of tools for DE analysis. First, limma-trend and voom were top performers for scRNA-Seq data demonstrated by a recent benchmarking study^14^. Secondly, the linear models of limma enabled complex design. Additionally, we benchmarked limma with other top performers (MAST^23^ and edgeR^24^) in a simulated dataset confounded by batch effects. MAST uses a hurdle model of two-part generalized linear model, aiming to model the bimodality expression pattern of zero-inflated scRNA-Seq data, whilst edgeR fits the coefficients and the dispersion parameters using negative binomial distribution. In our benchmarking experiment, the limma methods detected the signal-to-noise ratio as well as edgeR and better than MAST, and its computing speed was much faster (Supplementary Fig. 1a, b), which is consistent with previous report^12^. Moreover, limma-trend was faster than voom, because voom has an additional step of inferring variance at the level of individual observations. Our workflow provides both limma-trend and voom for choice. Limma-trend is recommended when the runtime is a major concern, while voom may perform slightly better when library sizes are unequal^12^.

IDER is aimed to measure the inter-group similarity. In the scenarios of multiple batches, IDER first compare two groups, *g_i_* and *g_j_*’, with the background, i.e., cells that do not belong to *g_i_* and *g_j_*’, respectively (Fig. 1 a). For each comparison, the DE analysis was performed with the linear regression including covariates of group *(g_i_, g_j_’*, and background), batch and scaled cellular detection rate. The cellular detection rate measures the number of genes detected per cell as previously described^23^. After the estimated coefficients are computed, the DE signature, vector *d_i_*, for group *g_i_* (or *d_j_*’ for group *g_j_’)* is computed by fitting the contrast of *g_i_* - background (or *g_j_*’ - background). The length of *d_i_* or *d_j_* is equal to the number of genes, i.e., number of rows in the count matrix. The DE signature can be either denoted by the estimated coefficients, i.e., log_2_ fold-change or sign(fold-change) * −log10(p), where p is the Benjamini-Hochberg-adjusted p values estimated by the moderate *t*-test.

Between the two groups, *g_i_* and *g_j_*’ the similarity *s(g_i_, g_j_’)* is measured by the consistency of DE signatures, *d_i_* and *d_j_’.* In this study, we used the Pearson correlation coefficients. Alternative options are Spearman, Kendall correlation coefficients and cosine similarity. The similarity measure ranges from −1 to 1.

IDER can also be used to measure inter-group similarity for data having confounding factors other than the batch, e.g., treatment. In this scenario, the additional covariates can be added into the regression. On the other hand, when there is only one batch, the design matrix can be simplified by removing the batch.

### CIDER for identifying cell populations

To cluster multi-batch data, CIDER consists of three steps: initial clustering, computing similarity matrix and final clustering. For dnCIDER, we first used Louvain clustering to cluster cells within each batch. Pairs of batch-specific clusters with high similarity of IDER were merged, generating the initial clusters for the next step. For asCIDER, we concatenated the batch tag and the cell annotation as the initial cluster. Next, to generate the similarity matrix, the pairwise similarity was computed for all cross-batch pairs of initial clusters by IDER. We downsampled each initial cluster to the same size (35 to 50 cells). We do not suggest downsampling to a number smaller than 15. For initial clusters smaller than this size, we allowed replacement for sampling. To visualize the similarity among initial clusters, this similarity matrix was transferred to a graph by using igraph in R (https://igraph.org/r/). In the final clustering step, the similarity matrix *S* was transferred to a distance matrix by (1 -***S***)/2 and the initial clusters were merged by the hierarchical agglomerative clustering with complete linkage, enabling the initial clusters with the highest similarity to be merged first. For large datasets, parallel computation (R package doParallel) was used to shorten the runtime.

### External data

#### Cell line data (Dataset 1)

We obtained the data of 293T cells and Jurkat cells from http://scanorama.csail.mit.edu/data.tar.gz^7,15^. This dataset came from three batches. The first batch has only 293T cells, the second batch only Jurkat cells, and the third batch 1:1 mixture of these two cell lines.

#### Human PBMC data (Dataset 3)

The raw count matrix and the sample information were downloaded from https://hub.docker.com/r/jinmiaochenlab/batch-effect-removal-benchmarking^10^, which were curated in the recent benchmarking study^10^. Cells with at least 500 genes detected were kept for further analysis. Putative doublets were filtered by DoubletFinder^25^ for each batch.

#### Cross-species pancreas data (Dataset 4)

The count matrix and sample information were downloaded from NCBI GEO accession GSE84133^16^. We kept cells with minimum 500 genes detected for downstream analysis. Doublets were filtered by DoubletFinder^25^. The gene set shared by both human and mouse was kept for downstream analysis. The human gene *INS* were treated as the mouse gene *Ins1* as previously described^8^.

#### COVID-19 data (Dataset 5)

The 10x data were downloaded from GSE149689^17^ and the cell annotations were downloaded from https://junglab.wixsite.com/home/db-link.

#### Human dendritic data (Dataset 6)

The data were downloaded from https://hub.docker.com/r/jinmiaochenlab/batch-effect-removal-benchmarking^10^ and contain four cell populations (CD141, CD1C, DoubleNeg and pDC) from two batches^18^. The raw count matrix and the sample information were also downloaded from the curated set^10^. Cells with less than 500 genes detected were removed.

#### Mouse hematopoietic progenitor data (Dataset 7)^19,20^

The data were downloaded from the curated set^10^ and contain three cell populations, named CMP, GMP and MEP, sequenced by two platforms (MARS-seq and Smart-seq). CMP was recognized as the direct ancestry of GMP and MEP^19,20^.

### Integration pipelines

#### Seurat CCA

We used the recommended CCA correction pipeline of Seurat v3.1.5^8^. We first split objects by batches, followed by normalization and selection of top 1000 HVGS based on the relationship between mean and variance. The integration anchors were identified from the first 15 PCs. After integration, the first fifteen PCs of the MNN-correct matrix was used for Louvain clustering with resolution of 0.4.

#### MNN

We used scran v1.14.5 to identify HVGs, which were used as the input of fastMNN^6^ (batchelor v1.2.4). The first fifteen PCs of the MNN-correct matrix was used for Louvain clustering with resolution of 0.4.

#### Scanorama

We used Scanorama^7^ via reticulate v1.16 in R as suggested by the Scanorama repository (https://github.com/brianhie/scanorama). The first fifteen PCs of the Scanorama-correct matrix was used for Louvain clustering with resolution of 0.4. Due to the limitation of memory, the data were downsampled to 1:10 for Dataset 5 before applying Scanorama.

#### Harmony

We used the RunHarmony function of Harmony v1.0^9^ to perform integration and used the first 15 corrected-PCs as the input of Louvain clustering with resolution of 0.4.

### Clustering pipelines

#### Seurat Louvain clustering

We used the suggested pipeline of Seurat v3.1.5^5^. The top 2000 HVGs were used to compute PCs, while the first fourteen PCs were used for Louvain clustering with resolution of 0.4.

#### SC3

We used SC3 v1.14.0^3^. The number of clusters based on ground truth was given to the clustering function.

#### RaceID

We used the suggested pipeline of RaceID v0.1.9^4^, including filterdata, getfdata, compdist and clustexp. The number of clusters based on ground truth was given to the clustering function. As the Scater object that SC3 and RaceID depended on consumed a substantial amount of memory, the data were downsampled to 1:10 for Datasets 3, 4 and 5 before applying SC3 and RaceID.

### Proof-of-concept analysis

The cell line dataset (Dataset 1) was corrected by Scanorama as previous described^7^. The first two components of t-SNE were used to perform Hierarchical DBSCAN (R package dbscan v1.1) with the minimum size of clusters set at 75. The output of DBSCAN and the batch information were combined to generate initial clusters. The Scanorama correction was used here as the ground truth, which has been demonstrated previously^7^. The initial clusters were downsampled to the size of 50 cells. The similarity matrix based on IDER was computed among the initial clusters to demonstrate the ability of capturing biological variance.

### Data simulation

We used Splatter v1.10.0^26^ to simulate scRNA-seq data. We first simulated a dataset with five groups and three batches and removed groups 4 and 5 from batch 1, groups 1 and 5 from batch 2 and groups 1 and 3 from batch 3. This generated the non-overlapped scenario (Dataset 2). The replications were generated in the same way with various seeds.

### Benchmarking clustering performance

The adjusted Rand index (ARI) was used to measure the consistency between clustering results and ground truth. We calculated the ARI between clustering results and the annotation of cell populations, termed ARI_population_. It indicates the accuracy of identifying cell populations. We also computed the ARI between clustering outcome and the annotation of batches, termed ARI_batch_. It represents the confounding effects of batches. Therefore, a larger value of 1-ARI_batch_ indicates that the clustering result is less confounded by batch effects. The runtime was tested on macOS with 3.2 GHz processor and 16 GB memory. To avoid overlong runtime or collapse of R, we downsampled the data for SC3 and RaceID by a factor stated in the figure legends.

### CIDER for evaluating integration

In this evaluation workflow, the batch-corrected low-dimensional representation was first used to partition cells into multi-batch clusters. These multi-batch clusters were further divided into batch-specific subclusters. Within each cluster, inter-group similarity was calculated between subclusters from a pair of batches, whilst the batch effects were regressed out by using the IDER metric. Higher levels inter-group similarity indicated better quality of integration for the cluster. For two batch-specific subclusters from the same cluster, we could estimate the probability whether they come from a true biological population (either cell type, subtype or state). To estimate the probability, we assumed that the two mutual nearest batch-specific groups with the highest similarity are from the same population (“mutual nearest neighbor” hypothesis) and that the variability within a given biological population is at an almost constant level (“constant variability” hypothesis). By further partitioning the combination of these two batch-specific subclusters, we can get a distribution of the variability within this merged cluster. An empirical p-value was next calculated for each pair of subclusters from the same cluster to indicate the probability of belonging to the same population. The LISI metric was used to validate the similarity calculated by CIDER. LISI_group_ was calculated as the LISI for the ground-truth annotations of cell populations in the batch-corrected t-SNE space. It was computed by using R package lisi^9^.

## Code availability

Analysis scripts of this work are deposited to the GitHub (https://github.com/zhiyhu/CIDER-paper).

## Acknowledgements

We thank Professor Eui-Cheol Shin for kindly providing the data. CY is supported by a UKRI-EPSRC Turing AI Fellowship (EP/V023233/1) and the UK Medical Research Council (MR/P02646X/2).

## Author contributions

ZH and CY conceived of the idea and designed experiments. ZH implemented the software and performed experiments. ZH, AAA and CY all contributed to the writing of the paper.

