## Supplementary Figures and Table for "An interpretable meta-clustering framework for single-cell RNA-Seq data integration and evaluation"

**a**

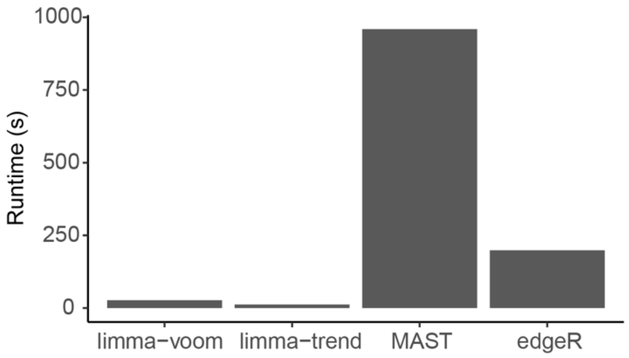


**b**

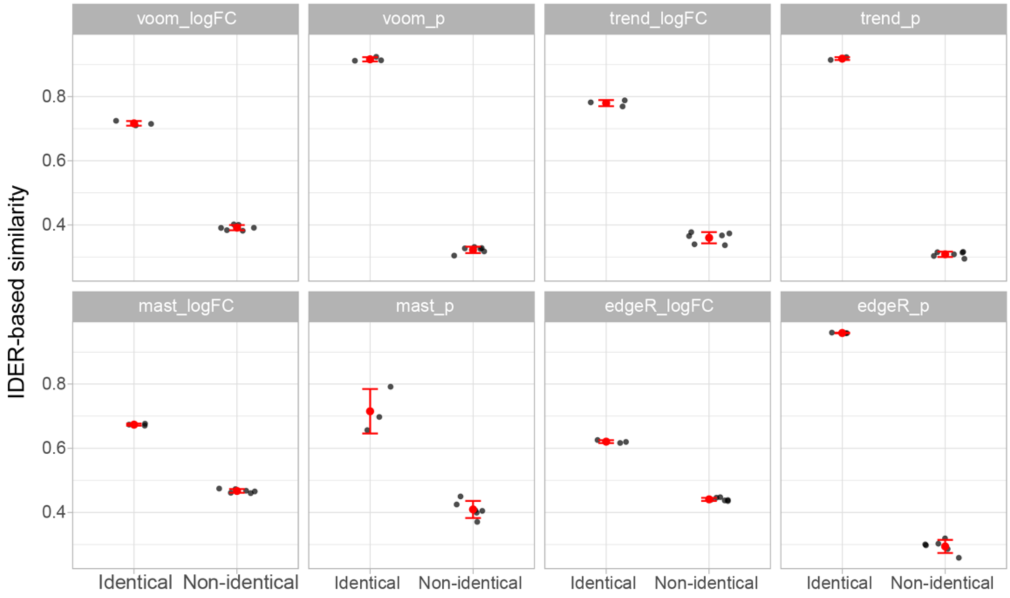


**Supplementary Fig 1. Benchmarking limma methods with MAST and edgeR for differential expression analysis.**

**(a)** Runtime of four methods used to compute the IDER matrix of a simulated dataset with two batches and 300 cells per batch. The computing speed is limma methods is faster than MAST and edgeR. Furthermore, limma-trend is faster than voom, because voom calculates observation-level variance in addition. Our workflow provides both limma-voom and limma-trend for choice, but limma-trend is recommended when the runtime is a major concern.

**(b)** The similarity levels computed by different methods. The limma methods detected the signal-to-noise ratio as well as edgeR and better than MAST. The x-axis shows the group pairs come from the identical population or non-identical populations. The y-axis denotes the IDER-based similarity between a pair of groups. Each dot is a group pair. The large red dot indicates median and error bars show standard deviations.

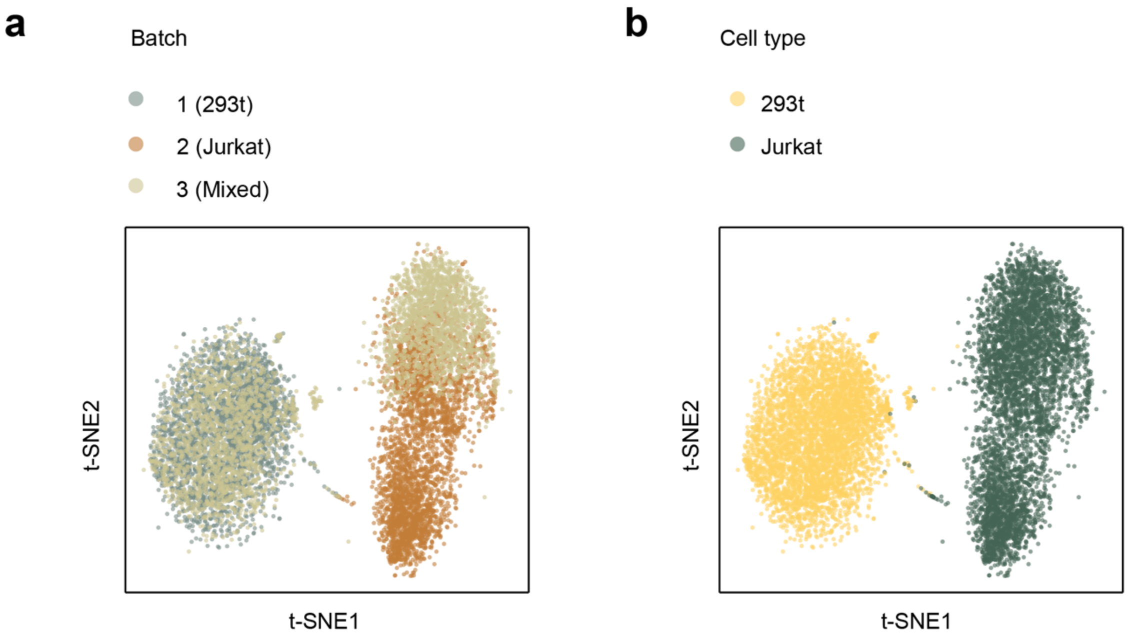


**Supplementary Fig 2. Compositions of the cell line dataset (Dataset 1).** (a-b) t-SNE plots of the cell line dataset corrected by Scanorama. Each dot denotes one cell, colored by the batch (a) and the cell type (b).

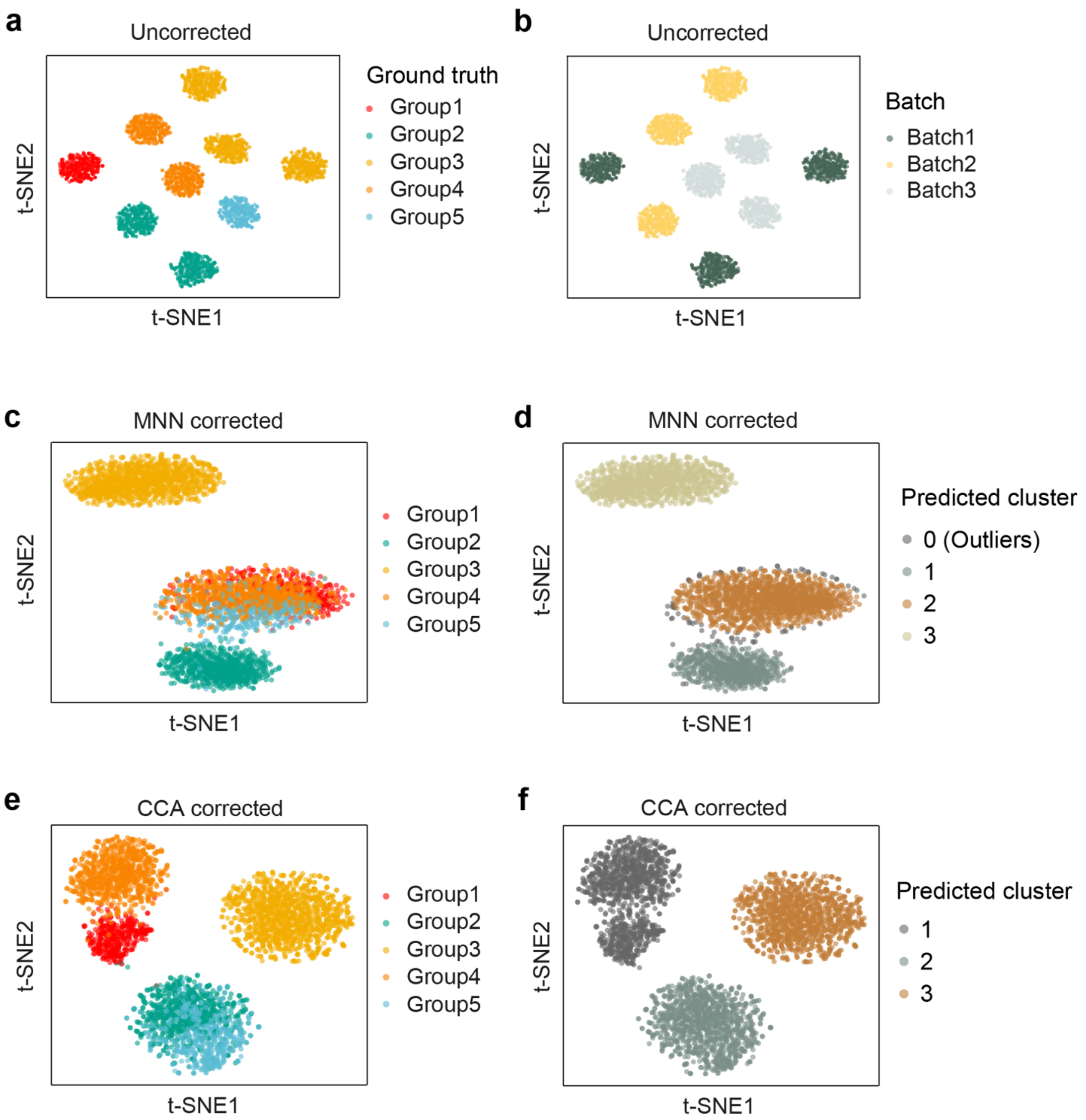


**Supplementary Fig 3. Compositions of the simulation dataset (Dataset 2). (a-b)** t-SNE plots of uncorrected data, where cells are colored by populations (a) and batches (b). **(c-d)** t-SNE plots of data corrected by MNN. Cells are colored by populations (c) and by clustering results of DBSCAN (d). **(e-f)** t-SNE plots of data corrected by Seurat-CCA. Cells are colored by populations (e) and by clustering results of DBSCAN (f).

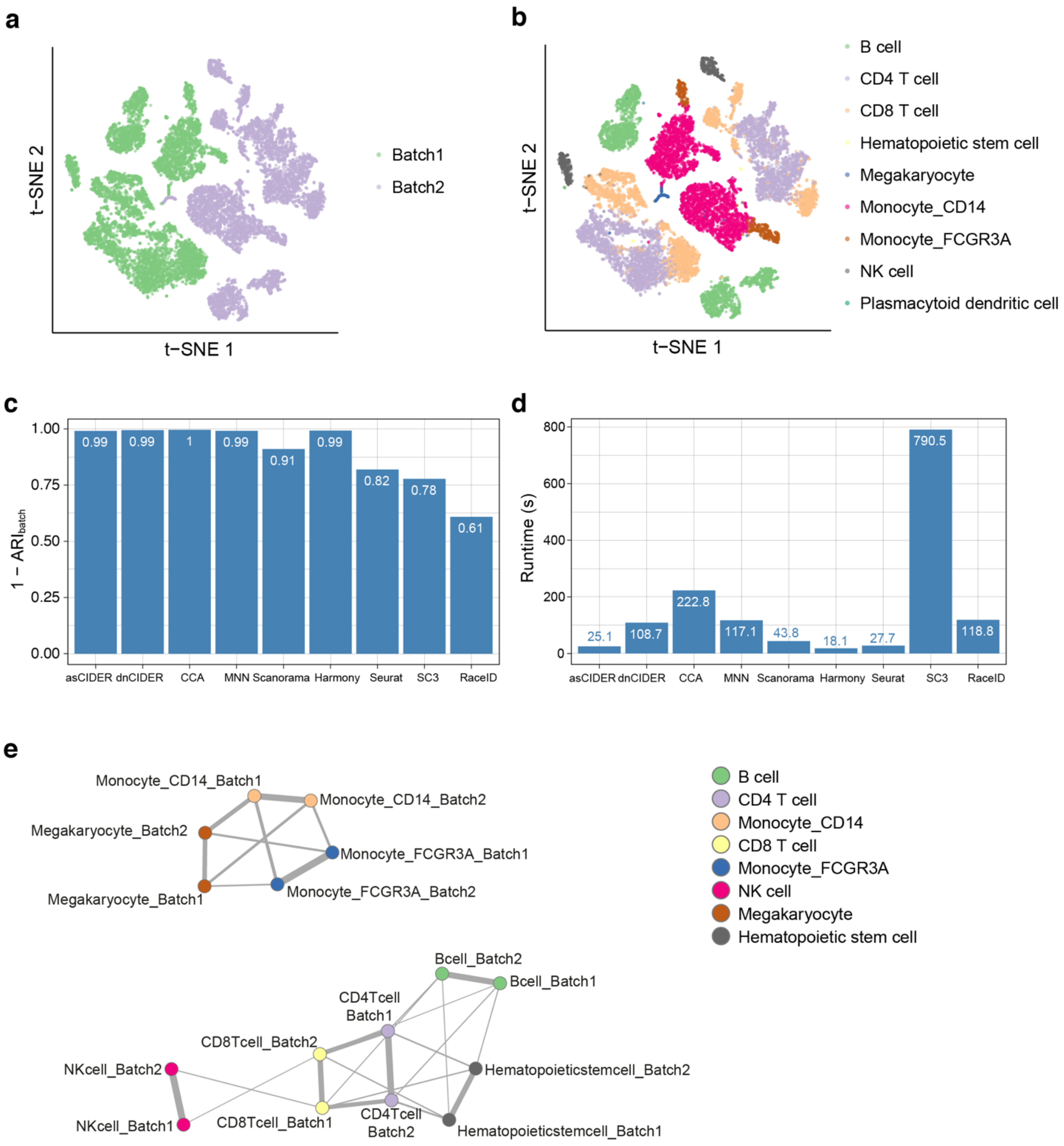


**Supplementary Fig 4. Benchmarking on the human PBMC data (Dataset 3). (a-b)** t-SNE plots of uncorrected Dataset 3, where cells are colored by batches (a) and populations (b). **(c)** Batch effects on clustering results by multiple workflows. The mere clustering workflows are more affected by batches. SC3 and RaceID were ran on the dataset downsampled by the factor of 10. **(d)** Runtimes of different workflows. SC3 and RaceID were ran on the dataset down sampled by the factor of 10. **(e)** Network shows the inter-group similarity among initial clusters.

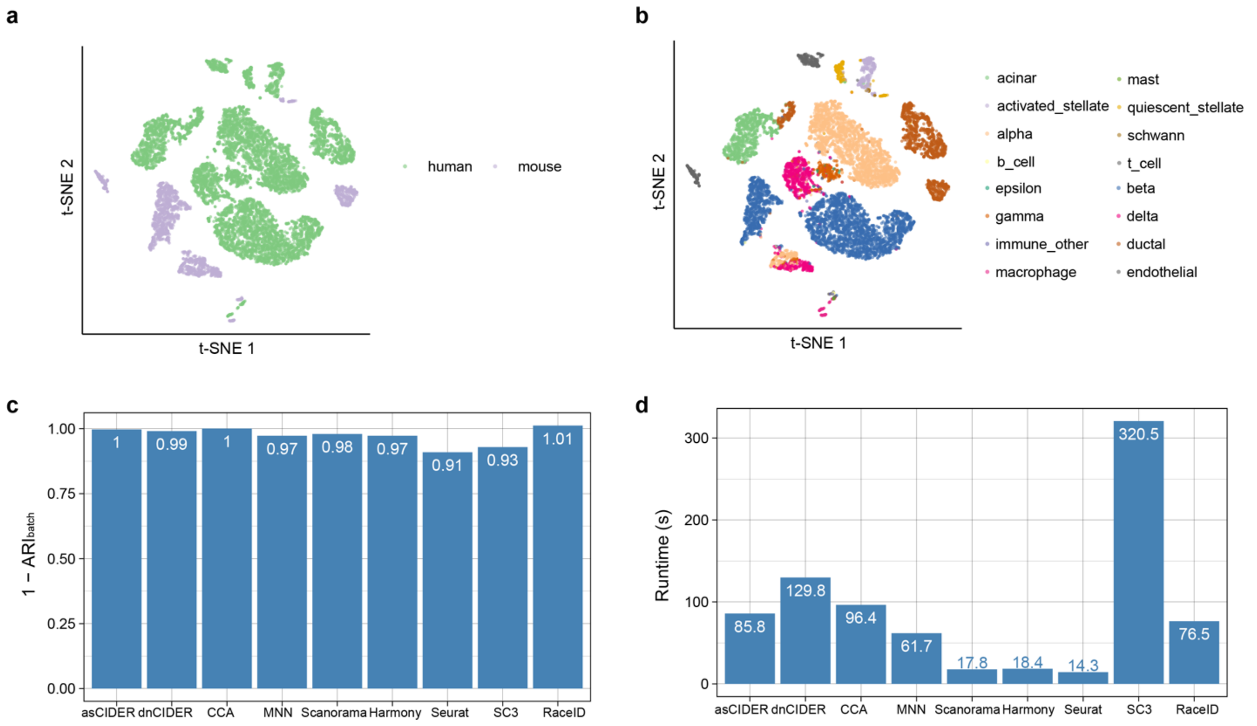


**Supplementary Fig 5. Benchmarking on the cross-species pancreas data (Dataset 4). (a-b)** t-SNE plots of uncorrected data, where cells are colored by batches (a) and populations (b). **(c)** Batch effects on clustering results by multiple workflows. **(d)** Runtimes of different workflows. SC3 and RaceID were ran on the dataset downsampled by the factor of 10.

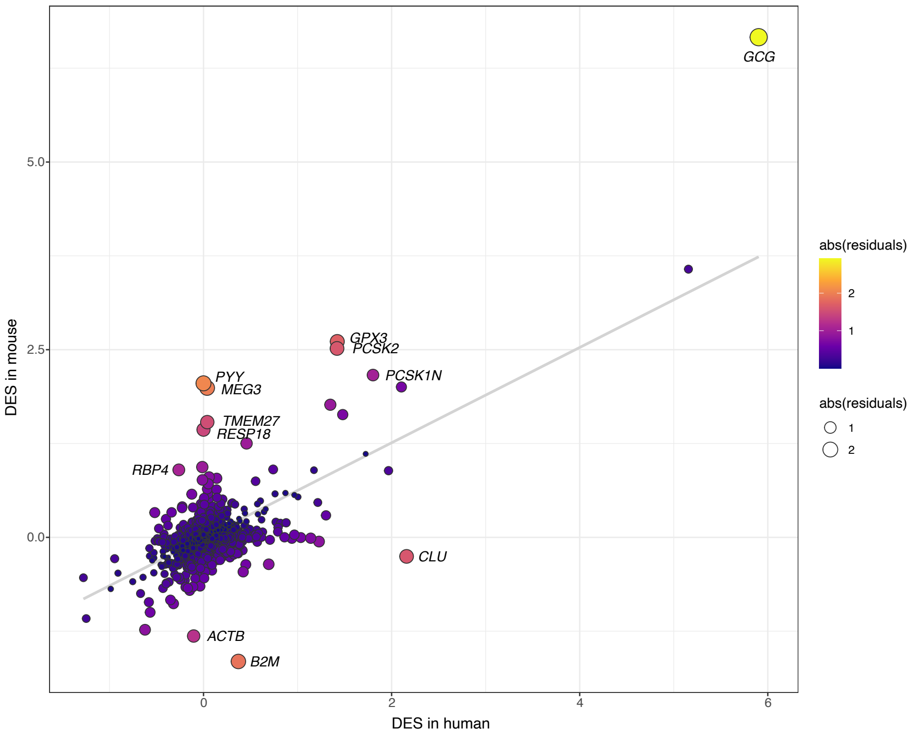


**Supplementary Fig 6. Biological interpretability of similarity between human and mouse alpha cells.** Scatter plot shows genes driving the dissimilarity between human ductal cells and mouse ones. The x- and y-axes denote the DESs in human and mice. Each dot is a gene, colored and sized by the abstract of residuals. The grey line is the linear regression line.

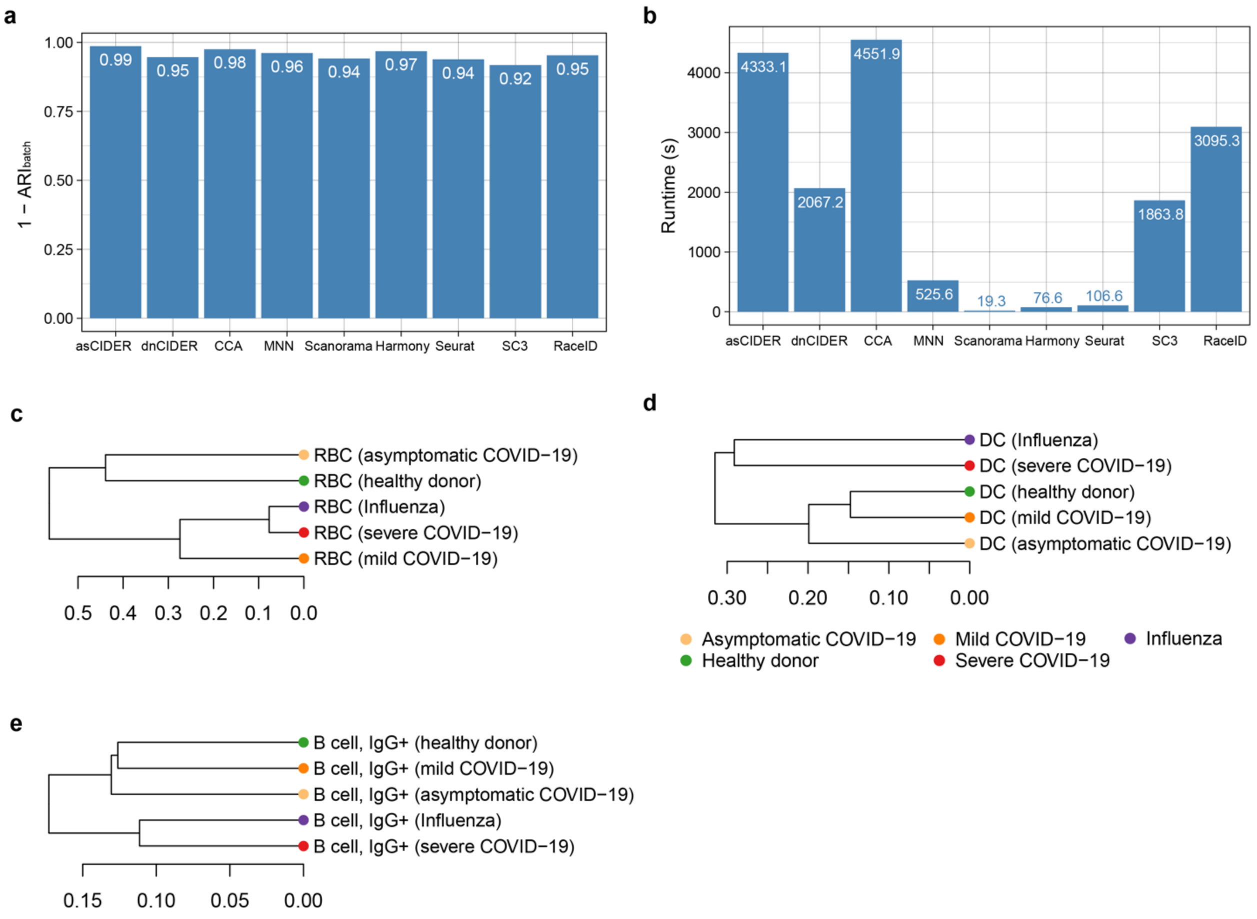


**Supplementary Fig 7. Benchmarking on the human PBMC data from patients with COVID-19, Influenza and healthy conditions. (Dataset 5). (a)** Batch effects on clustering results by multiple workflows. **(b)** Runtimes of different workflows. Scanorama, SC3 and RaceID were ran on the dataset downsampled by the factor of 10. **(c-d)** Dendrograms show the local relationships of the red blood cell (RBC) population (c), the dendritic cell (DC) population (d) and the IgG+ B cell population (e). Each leaf represents a cell population from one group of donors. The x-axis shows the height of branching.

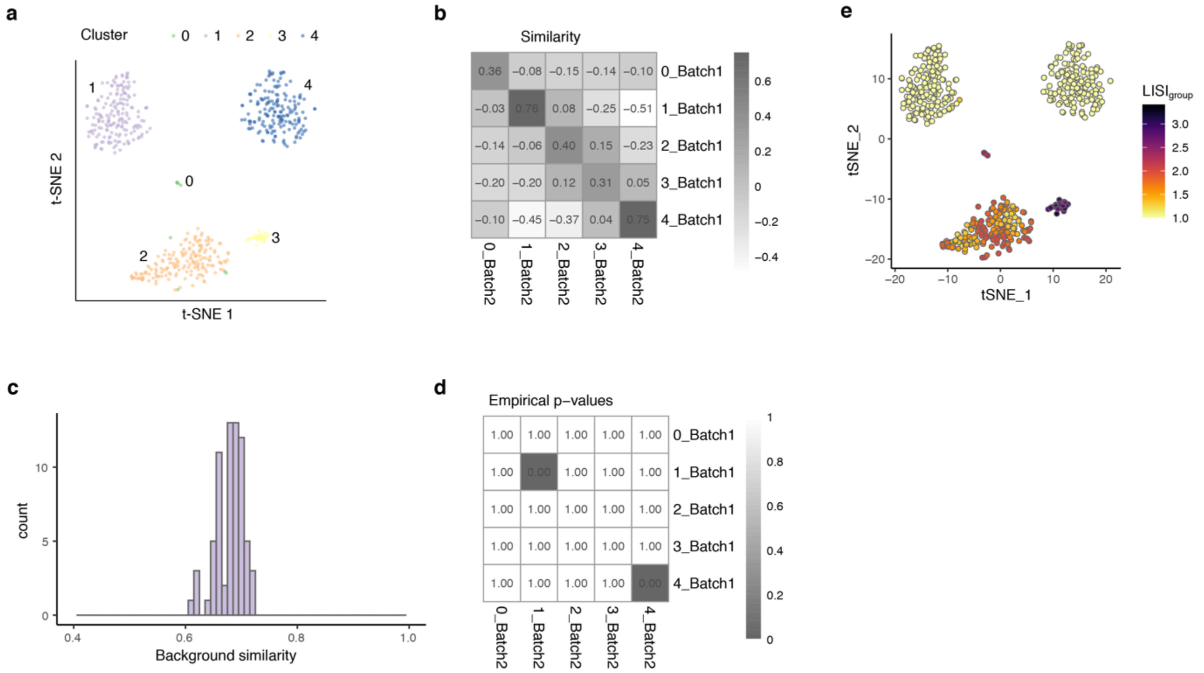


**Supplementary Fig 8. CIDER evaluates integration results by identifying mis-aligned population. (a)** t-SNE plot shows DBSCAN results of CCA-corrected dendritic data. **(b)** The IDER similarity between overlapped clusters from two batches. **(c)** The distribution of IDER similarity within the positive control groups that have the highest similarity. Here 1_Batch1 and 1_Batch2 are used. **(d)** The empirical p-value of whether the integration is acceptable computed by using the background distribution and the IDER matrix. **(e)** Evaluation by LISI requires the ground truth of cell populations. The t-SNE plot shows cells colored by $LISI_{group}$, a local diversity measure. A $LISI_{group}$ of 1 indicates identity purity of neighbors of a given cell.

**Supplementary Table 1. Summarized information of datasets.**

| ID | Data type | Cell numbers | Features |
|---|---|---|---|
| Dataset 1 | Cell line | 9530 | Non-overlap |
| Dataset 2 | Simulation | 6000 | Non-overlap |
| Dataset 3 | PBMC | 14876 | Different platforms |
| Dataset 4 | Pancreas | 10127 | Cross species |
| Dataset 5 | PBMC from COVID-19, Influenza and control conditions | 59572 | Diseases |
| Dataset 6 | Dendritic | 564 | Non-overlap |
| Dataset 7 | Mouse hematopoietic progenitors | 1442 | Trajectory |
